# Neurogenin 2 and Neuronal Differentiation 1 control proper development of the chick trigeminal ganglion and its nerve branches

**DOI:** 10.1101/2022.08.31.506039

**Authors:** Parinaz Bina, Lisa A. Taneyhill

## Abstract

The trigeminal ganglion contains the cell bodies of sensory neurons comprising cranial nerve V, which relays information related to pain, touch, and temperature from the face and head to the brain. Like other cranial ganglia, the trigeminal ganglion is composed of neuronal derivatives of two critical embryonic cell types, neural crest and placode cells. Neurogenesis within the cranial ganglia is promoted by Neurogenin 2 (Neurog2), which is expressed in trigeminal placode cells and their neuronal derivatives and transcriptionally activates neuronal differentiation genes like *Neuronal Differentiation 1* (*NeuroD1*). Little is known, however, about the role of Neurog2 and NeuroD1 during chick trigeminal gangliogenesis. To address this, we depleted Neurog2 and NeuroD1 from trigeminal placode cells with morpholinos and demonstrated that Neurog2 and NeuroD1 influence trigeminal ganglion development. While knockdown of both Neurog2 and NeuroD1 affected innervation of the eye, Neurog2 and NeuroD1 had opposite effects on ophthalmic nerve branch organization. Taken together, our results highlight, for the first time, functional roles for Neurog2 and NeuroD1 during chick trigeminal gangliogenesis. These studies shed new light on the molecular mechanisms underlying trigeminal ganglion formation and may also provide insight into general cranial gangliogenesis and diseases of the peripheral nervous system.

## 1. Introduction

The trigeminal ganglion houses the cell bodies and supporting glia of sensory neurons comprising cranial nerve V. These neurons arise from neural crest cells and placode cells, and reciprocal interactions between these cell types are critical to assemble the ganglion [1–3], which possesses ophthalmic, maxillary, and mandibular nerve branches [4]. In support of this, prior studies demonstrated that ablation of chick neural crest cells led to the scattering of trigeminal placode cell-derived neurons and the formation of two disconnected ganglia, indicating the importance of neural crest cells as an aggregating center [1]. On the other hand, placodal neurons are fundamental for formation of neural crest-derived neurons in the trigeminal ganglion [5]. Extirpation of chick trigeminal placode cells resulted in the absence of either the ophthalmic or maxillomandibular branches, or sometimes both branches, pointing to a critical role for placode cells in proper trigeminal ganglion formation [1].

In chick embryos, ophthalmic and maxillomandibular placode cells differentiate to form trigeminal sensory neurons, with the former appearing first in the surface ectoderm (E1, Hamburger Hamilton (HH)8) and the latter detected about 36 hours later (E2.5, HH16) [6]. By E2.5-3 (HH16-17), trigeminal placode cell-derived neurons have already delaminated and migrated to the ganglionic anlage, where they intermix with neural crest cells to begin forming a condensed trigeminal ganglion [3]. This process involves the guidance of placodal neurons from the epithelium to the hindbrain by neural crest cell streams [7], which form “corridors” to define this path [8]. It is only later in development that neural crest cells will differentiate into neurons (E4, HH22-24) [3] to generate the remaining sensory neurons in cranial nerve V.

Vertebrate neurogenesis is controlled, in part, by proneural genes encoding basic Helix-Loop-Helix (bHLH) transcription factors [9] that also regulate cell type determination and terminal differentiation [10]. One family of bHLH proteins are the Neurogenins (Neurogs), which consists of Neurogs1-3 [11]. In the chick, *Neurog1* is observed in the maxillomandibular trigeminal placode, the vestibulo-acoustic otic vesicle, and the epibranchial placodes [4, 12]. *Neurog2* is a chick ophthalmic placode-specific marker until E2.5 (HH16) [4, 12], after which it is considered a marker for all placode-derived neurons since its expression is detected in other placodes at E2.5 and in the maxillomandibular neurons of the trigeminal ganglion at E3 (HH18) [4]. *Neurog2* is also expressed transiently in a subset of neural crest cells [13]. *Neurog3* is not expressed in the chick trigeminal ganglion but is detected in the developing retina and in some cells of the non-neural retinal pigment epithelium [14]. Interestingly, the converse expression pattern is observed for mouse *Neurog1* and *Neurog2*, with *Neurog1* noted in the trigeminal and vestibulo-acoustic placodes [15], while *Neurog2* is primarily expressed in the epibranchial placodes [16]. Additionally, Neurogs play a key role in activating downstream bHLH factors such as *NeuroD1*, which is expressed in chick trigeminal placode cells prior to delamination and in their neuronal derivatives up to E8 [17]. Notably, Neurogs also regulate downstream signaling pathways controlling the neuronal cytoskeleton and subsequent morphology.

Despite our understanding of *Neurog2* and *NeuroD1* expression, little is known about Neurog2 and NeuroD1 function in the chick embryo, particularly with respect to the trigeminal ganglion. To this end, we depleted Neurog2 or NeuroD1 from chick trigeminal placode cells using validated morpholino antisense oligonucleotides (MOs) and evaluated trigeminal ganglion development. Our studies reveal, for the first time, a role for Neurog2 and NeuroD1 in chick trigeminal placode cells, and have shed light on mechanisms underlying trigeminal ganglion development.

## 2. Materials and Methods

### 2.1 Chicken embryos

Fertilized chicken eggs (*Gallus gallus*) were obtained from Centurion Poultry, Inc. (Lexington, GA) and the Department of Animal and Avian Sciences at the University of Maryland (College Park, MD) and incubated at 37°C in humidified incubators. After approximately 38 hours of incubation, eggs were removed from the incubator and a window was made in the shell to access the embryo. Staging was conducted according to the HH staging table [18]. Manipulations were performed on embryos at approximately E1.5 between the 8 somite stage (ss) to 10ss (HH9+ through to HH10). Embryos at E2 (HH13) and older were subsequently collected for analyses.

### 2.2 Morpholino design and electroporation

A 3′ lissamine-tagged translation-blocking Neurog2 MO (5′-TCTCCGCCTTCACCGGCATCC-3′), NeuroD1 MO (5’-CGGTGACGGTCGCATAACCCCG-3’), and a standard scrambled control MO prepared by the manufacturer (5′-CCTCTTACCTCAGTTACAATTTATA-3′), were designed to target each transcript or serve as a control, respectively, according to the manufacturer’s criteria (Gene Tools, LLC, Philomath, OR). All MOs were used at a concentration of 500 μM as previously described [18]. As recommended by Gene Tools, the inverse complement of the MO sequence was compared with the chicken transcriptome using the NCBI Nucleotide BLAST tool to test the selected target for homologies with other transcripts. These results revealed that the designed MOs only base pair with *Neurog2* and *NeuroD1* transcripts and are not complementary to any other sequences. Immunoblotting was also performed to demonstrate evidence of Neurog2 and NeuroD1 knockdown.

For electroporation, each MO was overlayed in a unilateral fashion on top of the ectoderm of∼E1.5 (HH9+ to 10) chick embryos (prior to placode cell delamination) by fine glass needles. After the MO was introduced, platinum electrodes were placed vertically across the chick embryo to deliver three pulses of 9 V, each lasting 50 milliseconds, at intervals of 200 milliseconds, as described [19]. Eggs were re-sealed with tape and parafilm and incubation was then continued for∼18-24 hours until the embryos reached E2 (HH13-14). A Zeiss SteREO Discovery.V8 microscope and X-Cite Fluorescence illumination (series 120) was then used to screen the embryos *in ovo* for the presence of the red fluorescent signal that emanates from MO-positive cells in order to confirm that trigeminal placode cells had been electroporated. After screening, eggs with successfully electroporated embryos were re-sealed and re-incubated for the desired time period.

### 2.3 Immunoblotting

The knockdown efficiency of both Neurog2 and NeuroD1 MOs was evaluated by collecting and pooling electroporated trigeminal ganglia dissected from Neurog2-(n = 25), NeuroD1-(n = 17), and control MO-(n = 25, n = 22, respectively) treated embryos at E2.5-3 (HH16-18). Samples were rinsed in Ringer’s solution, centrifuged at 500 g for five minutes at 4°C, and then snap-frozen in liquid nitrogen. Cell pellets were lysed in lysis buffer (50 mM Tris-HCl pH 8.0, 100 mM NaCl, 0.5% IGEPAL CA-630, 1 mM EDTA) supplemented with cOmplete™ Mini Protease Inhibitor Cocktail (Millipore Sigma, cat# 04693124001) and 1 mM PMSF (Millipore Sigma, cat# 10837091001) for 30 minutes at 4°C with mixing every 10 minutes. Following centrifugation at >20,000 g for 15 minutes at 4°C, the clarified, solubilized protein fraction was collected, and protein concentration was calculated using a Bradford assay (Bradford reagent, Bio-Rad, cat# 5000205). Each sample (containing equivalent amounts of protein) was boiled at 95°C for five minutes in 1X reducing Laemmli sample buffer and then centrifuged at maximum g for five minutes at room temperature. The samples were then loaded onto a 14% SDS-PAGE gel, separated by electrophoresis, and subsequently transferred to a 0.2 μm PVDF membrane (Thermo Fisher Scientific, cat# IB24002). Membranes were incubated in blocking solution (1X Phosphate-Buffered Saline (PBS) + 0.1% Tween (PTW) + 5% dry milk) for one hour at room temperature and then incubated overnight at 4°C with the following primary antibodies diluted in blocking solution: Neurog2 (1:200, Santa Cruz Biotechnology, cat# sc-293430) and NeuroD1 (1:1000, LifeSpan BioSciences, cat# LS-C331294). Membranes were washed three times in PTW for 10 minutes each and then incubated with species- and isotype-specific horseradish peroxidase-conjugated secondary antibodies at 1:15,000 dilution (Neurog2: mouse IgG-HRP, Rockland, cat# 610-1302; NeuroD1: rabbit IgG-HRP, Rockland, cat# 611-1302) in blocking solution for one hour at room temperature. PTW washes were repeated three times for 10 minutes each, and chemiluminescent substrates (Thermo Fisher Scientific, Supersignal West Pico, cat# 34580, or Femto, cat# 34095), along with a ChemiDoc XRS system (Bio-Rad), were used for detection. The immunoblots were then stripped (Restore Plus Western Blot Stripping Buffer, Thermo Fisher Scientific, cat# 46430) for 15 minutes at 37°C and re-probed with a loading control antibody (anti-Beta-actin primary antibody (1:1,500, Santa Cruz Biotechnology, cat# sc-47778, for the Neurog2 blot) and anti-GAPDH primary antibody (1:10,000, Thermo Fisher Scientific, cat# MA5-15738) for the NeuroD1 blot), followed by the appropriate secondary antibody (mouse IgG-HRP, 1:15,000, Rockland, cat# 610-1302). Immunoblots were analyzed using Image Lab software (Bio-Rad) in order to determine band size and volume.

### 2.4 Whole-mount immunohistochemistry

Fixed embryos in 4% paraformaldehyde were rinsed and then submerged in blocking solution (1X PBS + 0.1% Triton X-100 (0.1% PBST) + 10% sheep serum) for two hours at room temperature. Afterwards, the embryos were rinsed three times in 0.1% PBST for 10 minutes each. Embryos were then incubated overnight at 4°C with fresh antibody dilution solution containing primary antibody (Anti-Beta III Tubulin (Tubb3), 1:300, Abcam, cat# ab78078) in 0.1% PBST + 5% sheep serum, with gentle shaking. Next, embryos were washed four times for 30 minutes each at room temperature with 0.1% PBST, then incubated in fresh dilution solution with secondary antibody (goat anti-mouse IgG_2a_ AlexaFluor 488, 1:250, Southern Biotech, cat# 1080-30) overnight at 4°C with gentle shaking. Embryos were washed four times for 30 minutes each at room temperature with 0.1% PBST. Embryos were cleared before imaging, as described below.

### 2.5 Fructose and urea solution (FRUIT) clearing

Following immunohistochemistry, embryos were cleared via FRUIT, which utilizes a cocktail of fructose and urea to achieve maximum transparency of tissue without deformation [20]. Embryos were incubated in a series of FRUIT buffer solutions containing 8M urea (Millipore Sigma, cat# U5378), 0.5% (v/v) *α*-thioglycerol (Fisher Scientific, cat# T090525G), and increasing amounts of fructose (Millipore Sigma, cat# F3510). Embryos were gently rocked at room temperature in 35% FRUIT for six hours, 40% FRUIT for eight hours, 60% FRUIT for eight hours, and 80% FRUIT overnight. Embryos were kept at 4°C in 80% FRUIT before imaging.

### 2.6 Confocal imaging

Embryos were imaged in 80% FRUIT buffer on a Zeiss LSM 800 confocal microscope and Z-stacks were collected using 5X or 10X air objectives. When using contralateral control versus electroporated sides to image comparable regions of interest, the microscope laser power, gain, offset, and digital zoom were kept the same in each application, and the pinhole was always set to one airy scan unit. Zen software (Blue edition 2.0, Zeiss) was then used to process the CZI files. For Z-stacks, the CZI files were processed in ImageJ (NIH) [21], and the Z-Project function in HyperStack mode was used to create maximum intensity projections.

### 2.7 Measurement of ophthalmic branch width

The ophthalmic lobe width was measured on the electroporated and contralateral control sides of embryos treated with the Neurog2 MO (E3-3.5, HH18-20) and NeuroD1 MO (E2.5, HH16) using 5X and 10X maximum intensity Z-stack projections, respectively, at a distance of 100 µm from the point where the ophthalmic and maxillomandibular lobes separate. The measurement was conducted using the line tool in the open-source image processing program Fiji [22], which is based on ImageJ software [22]. A spatial calibration of the images was performed using Fiji based on the scale bar so that the measured distances measured are in microns.

### 2.8 Statistical analysis

Data associated with the width of the trigeminal ganglion ophthalmic branch are presented as boxplots. Boxes represent interquartile range, with the median value indicated as a line and whiskers representing the range. The Shapiro–Wilk test was used to assess distribution. Group differences were analyzed by the paired sample t-test. P-values equal to or below 0.05 were considered significant. All statistical analyses and boxplots were produced in R studio on R software (version 4.0.3) [23].

## 3. Results

### 3.1. Neurog2 controls the proper formation of the trigeminal ganglion and its nerve branches

To examine the function of Neurog2 during trigeminal ganglion neurodevelopment, Neurog2 knockdown experiments were carried out in trigeminal placode cells followed by immunohistochemistry on whole embryos to examine the forming trigeminal ganglion. To knockdown Neurog2 expression, a MO was designed to target the sequence surrounding the start site of the *Neurog2* transcript. The Neurog2 MO was unilaterally electroporated into the chick trigeminal placode ectoderm. Successfully electroporated embryos were re-incubated for specific periods of time, and then processed for further experimentation, as described below.

The efficacy of the Neurog2 MO was first tested by electroporating either a standard scrambled control MO (hereafter referred to as control MO) or the Neurog2 MO, followed by collection of electroporated trigeminal ganglia at E2-3 (HH16-18) to examine Neurog2 protein levels by immunoblotting [19]. Analysis of Neurog2 protein revealed a 30% reduction in the presence of the Neurog2 MO compared to the control MO (Figure 1).

**Figure 1.**
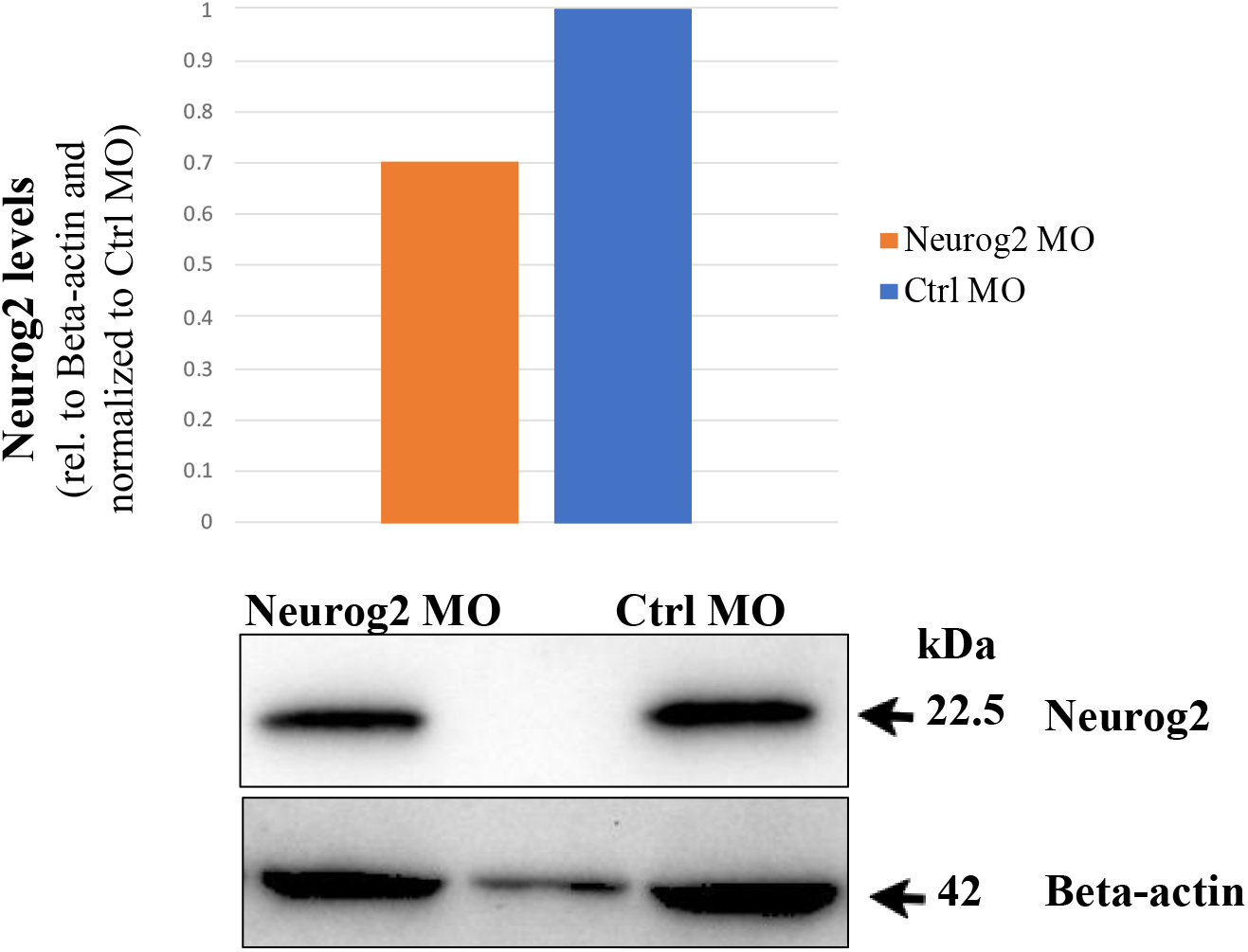
Neurog2 MO reduces Neurog2 protein levels by 30%. At ∼E1.5 (HH9+ to 10), placode cells were unilaterally electroporated either with a Neurog2 or control (Ctrl) MO. After re-incubation to E2.5-3 (HH16-18), the forming trigeminal ganglion on the electroporated side was dissected and pooled from multiple embryos. Lysates were prepared, and equivalent amounts of protein per sample were separated on a 14% SDS-PAGE gel. Immunoblotting for Neurog2 and Beta-actin (control) was then performed, and band intensity was calculated from unmodified immunoblot images using Image Lab software (Bio-Rad). Relative protein levels were ascertained by normalizing Neurog2 volumes to Beta-actin volumes. Knockdown amount was determined by comparing normalized ratios between Ctrl MO and Neurog2 MO samples, with the Ctrl MO sample set as one. On the Beta-actin panel, an extra band is present corresponding to lysate spillover from adjacent lanes.

To assess effects on trigeminal ganglion formation, successfully electroporated embryos were re-incubated to E3-5 (HH18-26), collected, fixed, and processed for whole-mount immunohistochemistry to detect Beta III tubulin (Tubb3), which labels differentiated neurons in the developing ganglion. At E2.5-3.5 (HH16-21), neuronal differentiation is occurring in the placodal population as neural crest cells will not begin differentiating into neurons until E4 (HH22-24)[1–3]; therefore, only placode-derived neurons will be identified. Confocal images of whole embryo heads were obtained to examine gross trigeminal ganglion morphology on the electroporated and contralateral control side, which possessed no MO.

At E3 (HH18), drastic changes in the trigeminal ganglion following Neurog2 depletion were already apparent. In contrast to the trigeminal ganglion on the contralateral control side (Figure 2A), the entire trigeminal ganglion and associated nerve structures were diminished in size on the Neurog2-depleted side (Figure 2B), which possessed many MO-positive cells (Figure 2C, F, arrows). Moreover, fewer axons were present in the ophthalmic branch, resulting in improper innervation of the eye region (Figure 2B, arrowheads). In addition, knockdown of Neurog2 appeared to alter the ability of the maxillomandibular branch to separate into definitive maxillary and mandibular branches, as shown by neurons deviating from the established maxillary branch (Figure 2A, B, D, E, carets). Besides these observations, however, the general morphology of the ganglion appeared similar: a bilobed structure possessing ophthalmic and maxillomandibular lobes and branches.

**Figure 2.**
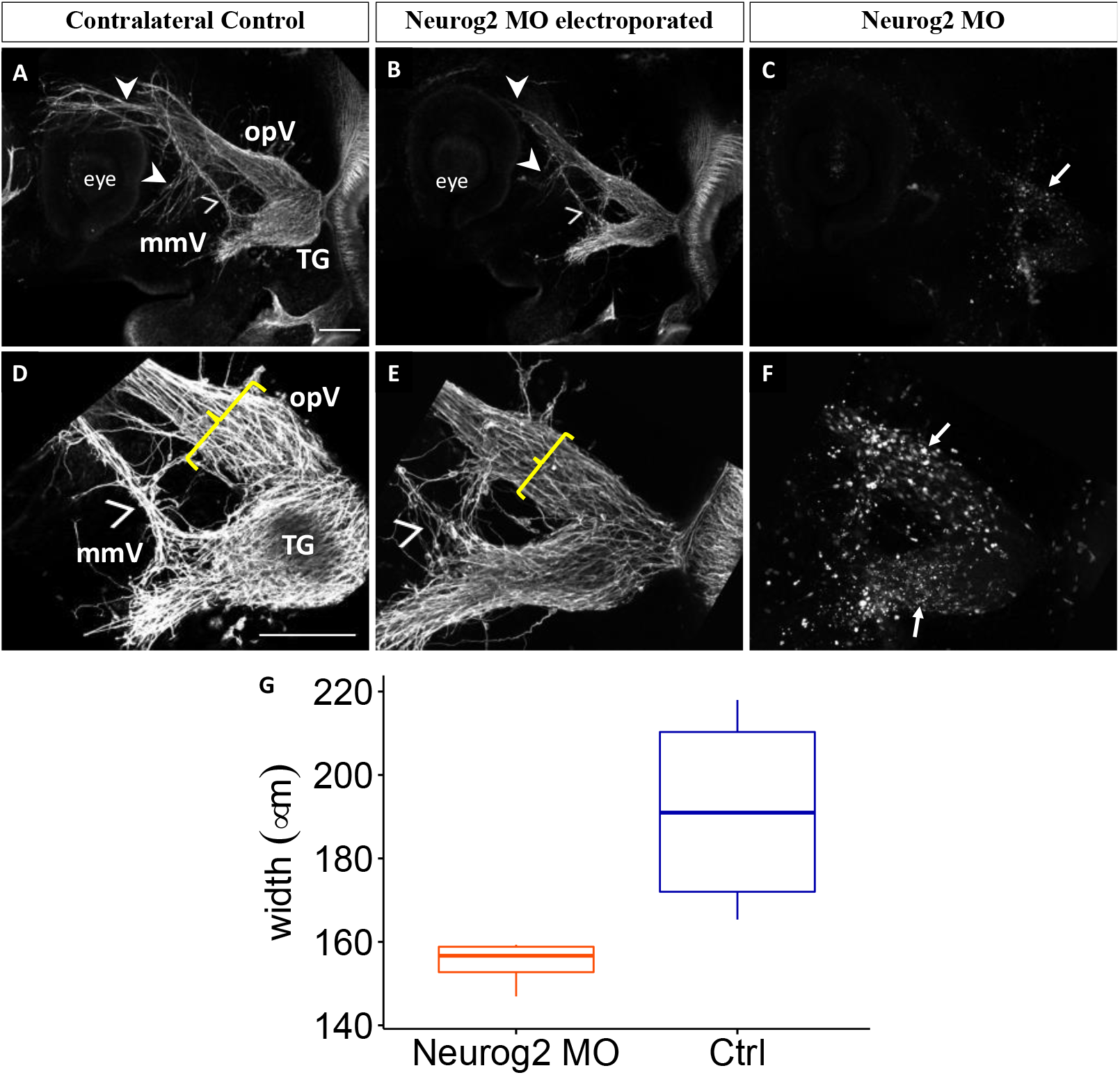
Depletion of Neurog2 in trigeminal placode cells impairs trigeminal ganglion development. Lateral view of the trigeminal ganglion in a chick head (E3 (HH18), n = 4). Representative images are maximum intensity projections of confocal Z-stacks through the contralateral control (A, D) and Neurog2 MO-electroporated (B, E) sides after processing for whole-mount Tubb3 Quantification of the width of the ophthalmic branch on the control (blue) and Neurog2 MO-treated (orange) sides. Plots represent the median (center line), 75th percentile (top of box), and 25th percentile (bottom of box), with whiskers connecting the largest and smallest values. A paired sample t-test revealed a p-value of 0.05. Scale bar is 1mm (A, D) and applies to all images. Abbreviations: mmV, maxillomandibular lobe; opV, ophthalmic lobe; and TG, trigeminal ganglion.

Although Tubb3-positive placodal neurons were observed throughout the forming ganglion, higher magnification images (Figure 2D-F) revealed neurons that were less organized and seemed to drift away from established axon bundles upon Neurog2 knockdown. Axons of the maxillomandibular nerve traveled without direction from the established nerve on the Neurog2-depleted side (Figure 2E) compared to the contralateral control side (Figure 2D). Moreover, the ophthalmic nerve branch was smaller in width on the electroporated side compared to the control (Figure 2D, E, brackets). Quantification revealed that this size difference was in fact statistically significant (p = 0.05, Figure 2G).

At E3-3.5 (HH20), the trigeminal ganglion and associated nerve structures were still reduced in size after Neurog2 knockdown (Figure 3B) compared to the trigeminal ganglion on the untreated contralateral control side (Figure 3A), with many MO-positive cells scattered throughout the forming ganglion (Figure 3C, F, arrows).

**Figure 3.**
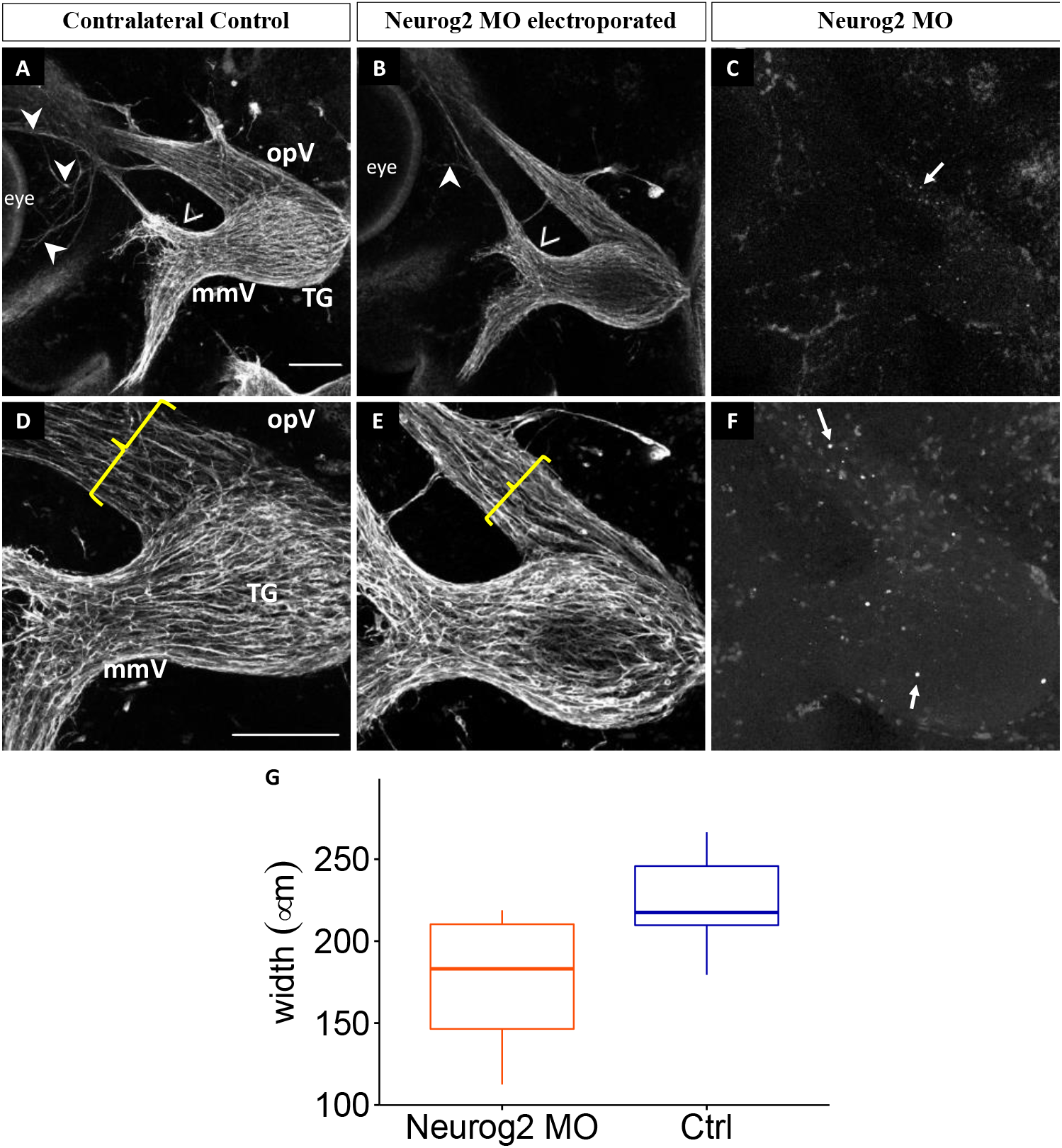
Neurog2 depletion in trigeminal placode cells disrupts trigeminal ganglion development. Lateral view of the trigeminal ganglion in a chick head (E3-3.5 (HH20), n = 6). Representative images are maximum intensity projections of confocal Z-stacks through the contralateral control (A, D) and Neurog2 MO-electroporated (B, E) sides after processing for whole-mount Tubb3 Quantification of the width of the ophthalmic branch on the control (blue) and Neurog2 MO-treated (orange) sides. Plots represent the median (center line), 75th percentile (top of box), and 25th percentile (bottom of box), with whiskers connecting the largest and smallest values. A paired sample t-test revealed a p-value of 0.02. Scale bar is 200μm (A, D) and applies to all images. Abbreviations: mmV, maxillomandibular lobe; opV, ophthalmic lobe; and TG, trigeminal ganglion.

Furthermore, the ophthalmic nerve extended less elaborately around the eye than on the contralateral control side (Figure 3A, B, arrowheads). Maxillary neurons were arranged in bundles but appeared less compact after Neurog2 knockdown than those on the untreated side (Figure 3A, B, carets). However, the trigeminal ganglion and its nerve branches appeared to have a similar overall morphology on the electroporated and contralateral control sides of examined embryos. With higher magnification (Figure 3D-F), though, a reduction in the width of the ophthalmic branch, and presence of likely fewer placode-derived neurons, were better appreciated (Figure 3D, E, brackets). This decrease in width was also statistically significant at this developmental stage (p = 0.02, Figure 3G). Collectively, these results reveal that Neurog2 knockdown impacts the development of the trigeminal ganglion and its nerve branches across multiple embryonic stages.

### 3.2. NeuroD1 regulates early chick trigeminal ganglion assembly

To further understand the function of bHLH factors in trigeminal ganglion development, we examined the role of NeuroD1 through knockdown experiments in trigeminal placode cells followed by immunohistochemistry on whole embryos. The NeuroD1 MO was designed to target the sequence surrounding the start site of the *NeuroD1* transcript. Successfully electroporated embryos were re-incubated to various developmental stages, and then processed for either immunoblotting or Tubb3 whole-mount immunohistochemistry, as described below.

We first tested the efficacy of the NeuroD1 MO by evaluating NeuroD1 protein levels through immunoblotting as we did previously for the Neurog2 MO. After electroporation, trigeminal ganglia at E2.5-3 (HH16-18) were then dissected and pooled for immunoblotting, which revealed three different bands immunoreactive with the NeuroD1 antibody, all of which are reduced in intensity after MO-mediated knockdown of NeuroD1 (Figure 4). Compared to NeuroD1 protein levels in the control MO sample, knockdown of NeuroD1 via the MO resulted in a 55%, 63%, and 31% decrease in the 50 kDa, 47 kDa, and 27 kDa NeuroD1 protein bands, respectively.

**Figure 4.**
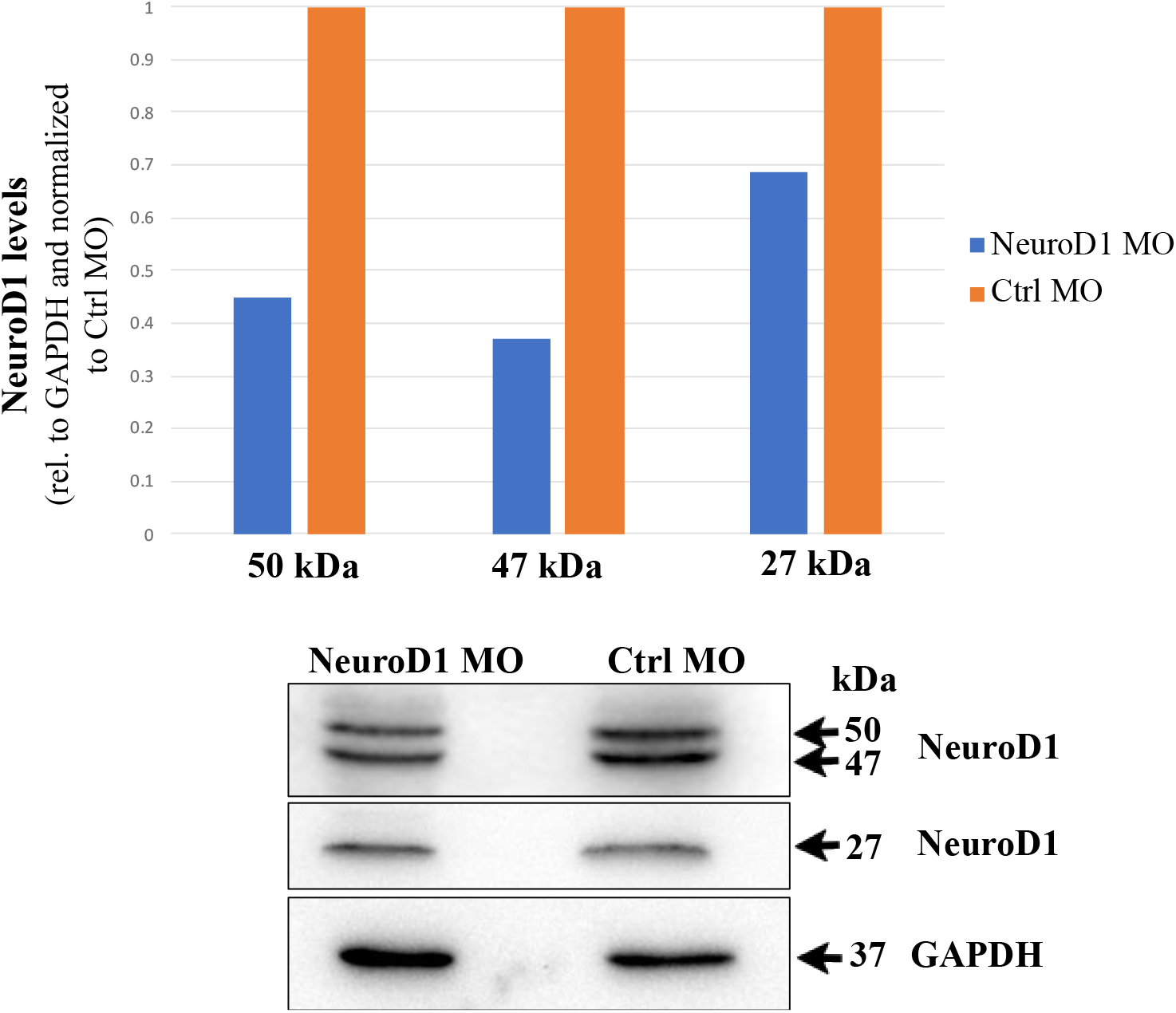
The NeuroD1 MO reduces NeuroD1 protein levels. At ∼E1.5 (HH9+ to 10), placode cells were unilaterally electroporated either with a NeuroD1 or control (Ctrl) MO. After re-incubation to E2.5-3 (HH16-18), the forming trigeminal ganglion on the electroporated side was dissected and pooled from multiple embryos. Lysates were prepared, and equivalent amounts of protein per sample were separated on a 14% SDS-PAGE gel. Immunoblotting for NeuroD1 and GAPDH (control) was then performed, and band intensity was calculated from unmodified immunoblot images using Image Lab software (Bio-Rad). Relative protein levels were ascertained by normalizing NeuroD1 volumes to GAPDH volumes. Knockdown amount was determined by comparing normalized ratios between Ctrl MO and NeuroD1 MO samples, with the Ctrl MO sample set as one.

To evaluate effects on trigeminal ganglion development, unilateral electroporation of NeuroD1 MO was conducted, followed by incubation of embryos to E2-2.5 (HH14-16), Tubb3 whole-mount immunohistochemistry, and confocal image acquisition, as carried out in our Neurog2 MO analyses. At E2 (HH14), changes in the trigeminal ganglion were already evident. In contrast to the trigeminal ganglion on the contralateral control side (Figure 5A, arrowhead), the forming trigeminal ganglion on the NeuroD1-depleted side was diminished in size and there appeared to be fewer neurons present (Figure 5B, arrowhead). Many MO-positive cells were also found in the electroporated ganglion (Figure 5C, F, arrows), and Tubb3-positive placodal neurons were observed throughout the condensing ganglion. Higher magnification images (Figure 5D-F) revealed neurons that were less organized. Accordingly, trigeminal sensory neurons on the electroporated side were widely dispersed (Figure 5E), whereas those on the control side were more densely packed (Figure 5D).

**Figure 5.**
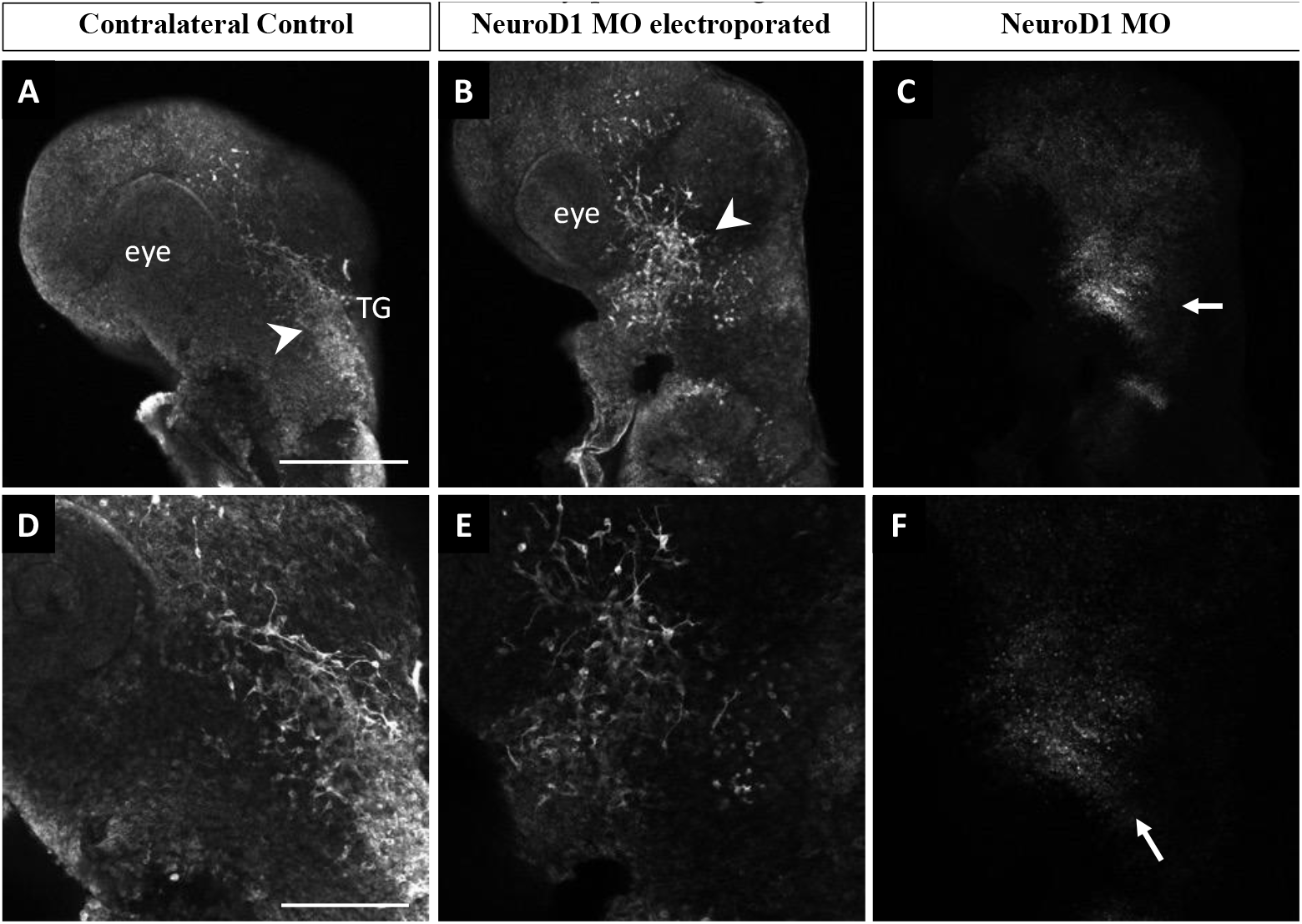
Depletion of NeuroD1 in trigeminal placode cells impairs trigeminal ganglion development. Lateral view of the trigeminal ganglion in a chick head (E2 (HH14), n = 3). Representative images are maximum intensity projections of confocal Z-stacks through the contralateral control (A, D) and NeuroD1 MO-electroporated (B, E) sides after processing for whole-mount Tubb3 immunohistochemistry to detect placode-derived neurons (A, B, D, E, arrowheads) and tissue clearing. Bottom row shows higher magnification images of the top row. (C, F) MO-positive cells (arrows). Scale bar is 500μm in (A) and applies to (B, C) and 200μm in (D) and applies to (E, F). Abbreviations: TG, trigeminal ganglion.

By E2.5 (HH16), both electroporated and control trigeminal ganglia possessed Tubb3-positive placodal neurons (Figure 6A, B, arrowheads), and MO-positive neurons were also observed in the electroporated ganglion (Figure 6C, F, arrows). However, there were differences in the way the neurites, and eventual axons, developed upon NeuroD1 knockdown (Figure 6A, B, arrowheads). Trigeminal sensory neurons on the contralateral control side extended axons into the eye area (Figure 6A, D), whereas those axons from NeuroD1 MO-electroporated trigeminal sensory neurons did not readily reach the eye (Figure 6B, E). Moreover, neurons within both lobes on the NeuroD1 MO-treated side exhibited an aberrant morphology (Figure 6E) compared to those on the contralateral side (Figure 6D). Additionally, trigeminal sensory neurons were more dispersed within the ophthalmic branch on the electroporated side compared to the control (Figure 6D, E, brackets). This increase in width upon NeuroD1 depletion was statistically significant at this developmental stage (p = 0.03, Figure 6G). Taken together, these results highlight a role for NeuroD1 in controlling the condensation of placodal neurons within the forming trigeminal ganglion.

**Figure 6.**
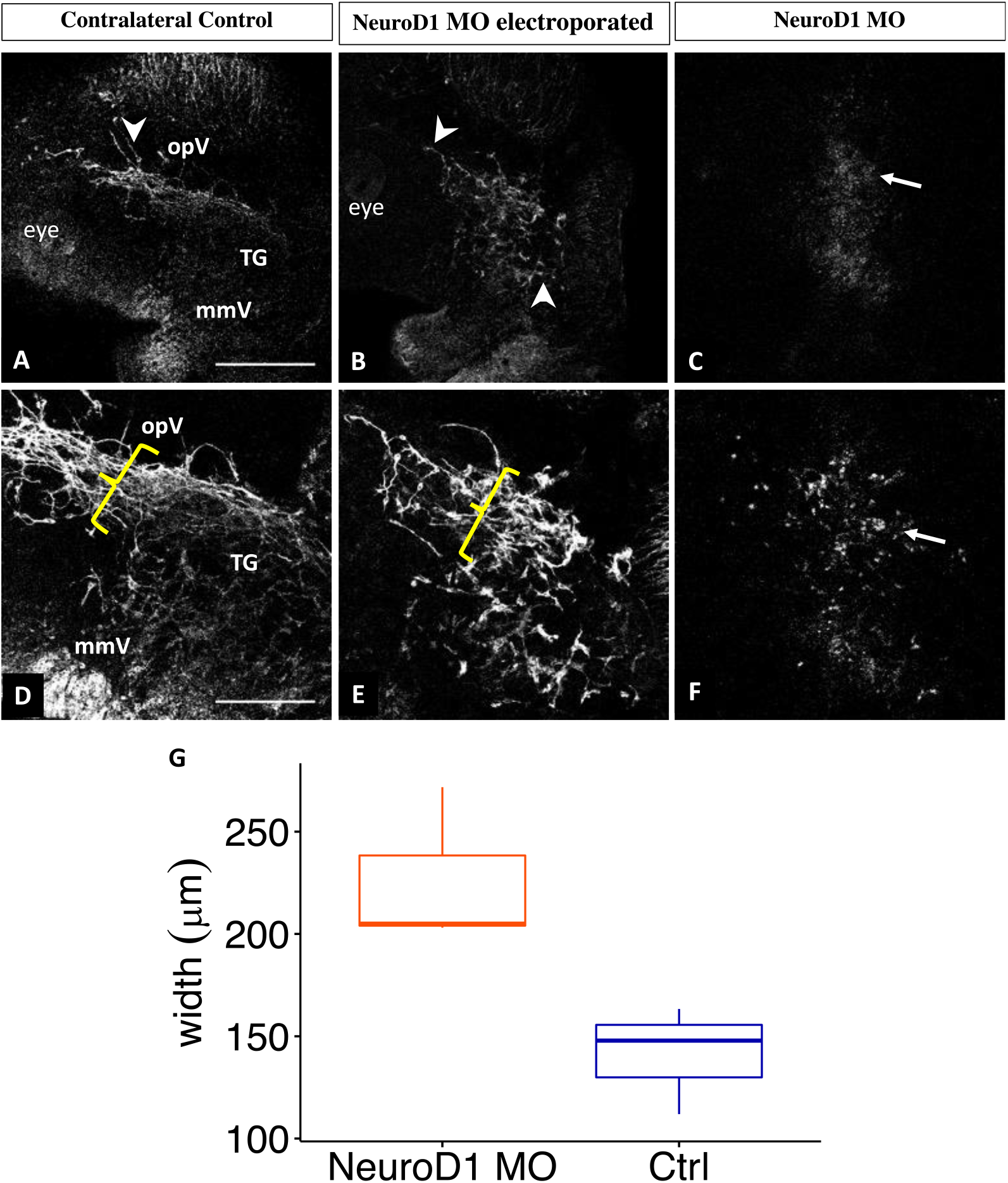
NeuroD1 depletion in trigeminal placode cells disrupts trigeminal ganglion development Lateral view of the trigeminal ganglion in a chick head (E2.5 (HH16), n = 3). Representative images are maximum intensity projections of confocal Z-stacks through the contralateral control (A, D) and NeuroD1 MO-electroporated (B, E) sides after processing for whole-mount Tubb3 immunohistochemistry to detect placode-derived neurons (A, B, D, E, arrowheads) and tissue clearing. Bottom row shows higher magnification images of the top row. (C, F) MO-positive cells (arrows). Brackets indicate width of ophthalmic branch. (G) Quantification of the width of the ophthalmic branch on the control (blue) and NeuroD1 MO-treated (orange) sides. Plots represent the median (center line), 75th percentile (top of box), and 25th percentile (bottom of box), with whiskers connecting the largest and smallest values. A paired sample t-test revealed a p-value of 0.03. Scale bar is 500μm in (A) and applies to (B, C) and 200μm in (D) and applies to (E, F) Abbreviations: mmV, maxillomandibular lobe; opV, ophthalmic lobe; and TG, trigeminal ganglion.

## 4. Discussion

Reciprocal interactions between neural crest cells and trigeminal placode cells are required to form the cranial trigeminal ganglion [1–3], which is involved in the perception of many sensations in the head and face, including touch, pressure, temperature, and pain [25].While this dual cellular origin of the trigeminal ganglion has been known for decades [1], the molecules mediating this process remain obscure. Neurogs belong to the bHLH transcription factor family and are known to play a crucial role in the development of placode-derived cranial sensory neurons. In the chick embryo, *Neurog2* is expressed primarily in ophthalmic trigeminal placodes, and is required to form trigeminal sensory neurons [4, 12]. Other transcription factors likely facilitate development of placode-derived sensory neurons by acting downstream of Neurogs [12]. *NeuroD1* has been suggested to be a target of Neurogs as revealed by *in situ* hybridization studies in mouse [15], but this has not been rigorously examined.

Although their expression pattern has been previously documented in the chick embryo, how Neurog2 and NeuroD1 function in the context of trigeminal placode cells and their neuronal derivatives is still poorly understood. To this end, we examined the role of Neurog2 and NeuroD1 during chick trigeminal gangliogenesis. Through knockdown experiments in chick trigeminal placode cells, we uncovered unique functions for Neurog2 and NeuroD1 in the forming trigeminal ganglion. Our results provide important insight into the role of these critical transcription factors in chick placodal neurons during trigeminal ganglion development.

### 4.1. Neurog2 regulates proper development of the trigeminal ganglion, and particularly the forming ophthalmic branch

To address the function of Neurog2 in chick trigeminal gangliogenesis, MO-mediated knockdown of Neurog2 was carried out in trigeminal placode cells. Despite achieving only 30% reduction in Neurog2 protein levels (Figure 1), Neurog2 MO treatment clearly caused dramatic effects on trigeminal gangliogenesis (Figures 2, 3), providing evidence that this protein is important for trigeminal ganglion development. Compared to the contralateral control side trigeminal ganglion, the ophthalmic branch of the trigeminal ganglion extended less elaborately around the eye on the Neurog2-depleted side of the embryo (Figure 2, 3). Further, Neurog2 knockdown resulted in a statistically significant decrease in the size of the ophthalmic branch compared to this branch of the trigeminal ganglion on the contralateral control side (Figure 2, 3-G). Additionally, Neurog2 depletion appeared to impair the segregation of the maxillomandibular branch into definitive maxillary and mandibular branches in the E3 trigeminal ganglion (HH18, Figure 2). Axons of the maxillomandibular branch on the Neurog2-depleted side also seemed less compact compared to those on the contralateral control side at E3-3.5 (HH20, Figure 3).

A reduction in trigeminal ganglion size could be due to, among other things, delayed delamination of placode cells contributing to the ganglion, increased cell death, or both. However, MO-positive cells are abundant in the ganglion, including in tissue sections (not shown), indicating that many electroporated cells have delaminated and migrated from the ectoderm. Thus, we can speculate that Neurog2 knockdown did not completely prevent placode cell delamination and migration. To determine if Neurog2 depletion delays delamination, live imaging of developing fluorescently-labeled placode cells could be carried out, which is beyond the scope of this study. Increased cell death could be ascertained over developmental time through TUNEL and/or immunohistochemistry to identify apoptotic cells within the forming ganglion. Although we hypothesize that the effects of Neurog2 (and NeuroD1) knockdown will be cell autonomous, future experiments should examine potential changes in the neural crest cell population to rule out any non-cell autonomous effects, particularly given the importance of neural crest-placode cell interactions as the trigeminal ganglion forms [1–3].

### 4.2. NeuroD1 influences trigeminal ganglion development

To ascertain the function of NeuroD1 in chick trigeminal ganglion development, MO-mediated knockdown of NeuroD1 was performed in trigeminal placode cells. Upon electroporation with NeuroD1 and control MOs, immunoblotting for NeuroD1 protein revealed three distinct bands (Figure 4). NeuroD1 protein has a predicted molecular weight of approximately 39 kDa in chick (Uniprot); however, immunoblot data has shown bands of various molecular weights (e.g., antibody websites), with a predominant band at 50 kDa [26–30]. Given that all three bands showed a reduction after knockdown, we conclude that all three bands represent NeuroD1 protein. The presence of bands at higher molecular weights than the predicted NeuroD1 protein product could be caused by post-translational modifications such as phosphorylation of NeuroD1, leading to a shift in electrophoretic mobility. This type of modification is not without precedence, as phosphorylation of Ser336 in NeuroD1 is essential for certain developmental processes, including dendrite growth and formation [31]. Additionally, NeuroD1 protein stability is regulated by ERK-dependent phosphorylation, which, in this instance, leads to ubiquitination and NeuroD1 degradation by the proteasome [26]. As such, the 27 kDa band could be a degradation product. Accordingly, a 55%, 63%, and 31% reduction in NeuroD1 protein levels impaired trigeminal gangliogenesis, indicating that this protein is critical for trigeminal ganglion development.

Depletion of NeuroD1 from trigeminal placode cells negatively affected trigeminal ganglion development. Axons from the ophthalmic branch of the trigeminal ganglion did not properly innervate the eye region and maxillomandibular neurons also possessed an abnormal morphology (Figure 6). Notably, ophthalmic and maxillomandibular neurons appeared dispersed and less compact after NeuroD1 knockdown compared to those on the contralateral control side of the embryo (Figure 5, 6), and these findings for the ophthalmic branch were statistically significant (Figure 6). As discussed in *Section 4.1*, these changes in the trigeminal ganglion and its nerve branches could be due to alterations in placode cell delamination and/or cell death, or potential effects on neural crest cells. As with Neurog2 MO treatment, we noted NeuroD1 MO-positive cells in the trigeminal ganglion in tissue sections (not shown), suggesting that effects on delamination are not necessarily substantial, at least at the stages we examined.

### 4.3. Possible roles for Neurog2 and NeuroD1 in trigeminal gangliogenesis

While our studies do not address the mechanism(s) by which Neurog2 and NeuroD1 control chick trigeminal gangliogenesis, prior work sheds some light on this, particularly with respect to effects noted on axon branching and neuron morphology. Studies in *Xenopus* demonstrated that Neurogs and NeuroD1 transcriptionally regulate genes whose protein products function in controlling the assembly and arrangement of cytoskeletal elements necessary for neuronal differentiation and migration [32]. Moreover, findings from the *Neurog2* knockout mouse identified the expression of cytoskeletal regulators to be negatively impacted [15]. Cytoskeletal changes are critical for neurons to make axons and dendrites from initially immature neurites, with rearrangements of actin filaments and microtubules dynamically occurring in neurites and in growing axons [33]. Thus, it is possible that placodal neuron morphology is affected due to intracellular changes occurring upon depletion of Neurog2 and/or NeuroD1.

Moreover, axon growth is regulated by guidance molecules, adhesion proteins, and neurotrophic factors [34]. The aberrant innervation of the eye that we observe after Neurog2 and NeuroD1 knockdown (Figures 2, 3, 6) could point to dysregulation of genes involved in these processes, such as those encoding neurotrophin receptors and/or neurotrophins, since lack of neurotrophic support leads to target innervation defects and neuronal cell death [29, 30]. Alternatively, it is possible that ophthalmic branch axons reach their target tissues normally after Neurog2 or NeuroD1 depletion, but are then retracted due to compromised cytoskeletal modifications caused by Neurog2 and/or NeuroD1 knockdown, as discussed above, which could be examined in the chick system in future experiments.

## 5. Conclusions

Our studies herein reveal that Neurog2 and NeuroD1 are critical to neurogenesis and the successful development of the trigeminal ganglion and its nerve branches. Through knockdown experiments, we demonstrated that Neurog2 and NeuroD1 are important for precise axon outgrowth and innervation of target tissues as well as neuron morphology. Altogether, our results provide new insight into molecules important for proper formation of trigeminal placode cell-derived neurons and will advance our understanding of trigeminal gangliogenesis in the chick embryo.

## Author Contributions

Conceptualization, P.B. and L.A.T.; methodology, P.B. and L.A.T.; validation, P.B. and L.A.T.; formal analysis, P.B. and L.A.T; investigation, P.B.; resources, L.A.T.; data curation, P.B.; writing—original draft preparation, P.B.; writing—review and editing, L.A.T.; visualization, P.B.; supervision, L.A.T.; project administration, L.A.T.; funding acquisition, L.A.T. All authors have read and agreed to the published version of the manuscript.

## Institutional Review Board Statement

Not applicable. At the stages of development being examined, the chick embryos used for our experiments are not considered live animals. According to the Office of Laboratory Animal Welfare (National Institutes for Health), “PHS Policy is applicable to proposed activities that involve live vertebrate animals. While embryonal stages of avian species develop vertebrae at a stage in their development prior to hatching, OPRR has interpreted “live vertebrate animal” to apply to avians (e.g., chick embryos) only after hatching.” Since our work does not utilize hatched chicks, no IACUC protocol for this work is necessary.

## Data Availability Statement

The data presented in this study are openly available in Digital Repository at the University of Maryland, Animal & Avian Sciences Research Works: Neurogenin 2 and Neuronal Differentiation 1 control proper development of the chick trigeminal ganglion and its nerve branches at [http://hdl.handle.net/1903/29101], reference number [37].

This project contains the following underlying data:

- Figure 1: Raw western blot data and excel file of band intensity quantification (Original raw tiff files for the immunoblotting experiments)
- Figure 2: Whole embryo images following electroporation of Neurog2 MO into placode cells (Original raw tiff files for the immunohistochemistry experiments)
- Figure 3: Whole embryo images following electroporation of Neurog2 MO into placode cells (Original raw tiff files for the immunohistochemistry experiments)
- Figure 4: Raw western blot data and excel file of band intensity quantification (Original raw tiff files for the immunoblotting experiments)
- Figure 5: Whole embryo images following electroporation of NeuroD1 MO into placode cells (Original raw tiff files for the immunohistochemistry experiments)
- Figure 6: Whole embryo images following electroporation of NeuroD1 MO into placode cells (Original raw tiff files for the immunohistochemistry experiments)

## Acknowledgments

We thank Ms. Claire Wegner for technical assistance.

